# Emotional arousal enhances narrative memories through functional integration of large-scale brain networks

**DOI:** 10.1101/2025.03.13.643125

**Authors:** Jadyn S. Park, Kruthi Gollapudi, Jin Ke, Matthias Nau, Ioannis Pappas, Yuan Chang Leong

## Abstract

Emotional events tend to be vividly remembered. While growing evidence suggests that emotions have their basis in brain-wide network interactions, it is unclear if and how these whole-brain dynamics contribute to memory encoding. We combined fMRI, graph theory, text analyses, and pupillometry in a naturalistic context where participants recalled complex narratives in their own words. Across three independent datasets, emotionally arousing moments during narrative perception were associated with an integrated brain state characterized by increased cohesion across functional modules, which in turn predicted the fidelity of subsequent recall. Network integration mediated the influence of emotional arousal on recall fidelity, with consistent within- and between-network interactions supporting the mediation across datasets. Together, these results suggest that emotional arousal enhances memory encoding via strengthening functional integration across brain networks. Our findings advance a cross-level understanding of emotional memories that bridges large-scale brain network dynamics, affective states and ongoing cognition.

---

Events that induce emotional arousal are often better remembered than those that are emotionally neutral^1–4^. For instance, a person is more likely to recall witnessing a car accident than what they saw on a routine drive home. Enhanced memory for arousing events prioritizes the recall of experiences that are motivationally significant, such as those relevant to potential threats and rewards^5–7^. This prioritization facilitates the efficient allocation of cognitive resources towards events that impact well-being, enabling individuals to respond more rapidly and effectively to similar events in the future^8,9^.

Emotional arousal is commonly defined as a subjective state of heightened alertness and activation^10–12^, often accompanied by physiological changes such as increased pupil dilation and electrodermal activity^13,14^. Prevailing theories propose that emotional arousal triggers the release of norepinephrine in the amygdala, which interacts with other memory-related regions, most prominently the hippocampus, to enhance the encoding of emotional experiences^15–18^. Consistent with this account, human functional magnetic resonance imaging (fMRI) studies have found that enhanced memory of emotional stimuli is associated with amygdala and hippocampal activation^19–23^, as well as increased amygdala-hippocampal connectivity^24,25^. These findings have been recently corroborated in intracranial recordings in patients^26,27^. The administration of the *β*-adrenergic antagonist propranolol, which interferes with norepinephrine binding, reduces amygdala and hippocampal responses to emotional stimuli and diminishes the associated memory enhancement, suggesting a causal role of arousal-dependent norepinephrine release in mediating these effects^28–31^.

Current models of arousal-dependent memory enhancement have made foundational contributions to our understanding of the brain regions involved, but they generally do not consider the role of the large-scale functional network organization of the brain (c.f. ^32^). There is growing recognition that the dynamic reconfiguration of distributed functional brain networks in response to task demands is crucial for adaptive behavior^33–38^ and underlies emotional responses and affective states^12,39–42^. Converging evidence suggests that heightened arousal promotes brain network integration - a brain state characterized by increased connectivity and cohesion across functional brain systems^33,43–48^, which have in turn been associated with memory encoding and retrieval^49,50^. Network integration is thought to be supported by the release of norepinephrine, as demonstrated by pharmacological manipulations^46,48^, chemogenetic stimulation^45^, and pupil dilation^33,43^, a widely used proxy of norepinephrine activity^51^. Building on this work, we propose that functional network integration facilitates the enhanced encoding of emotionally arousing memories, above and beyond the contribution of the amygdala and the hippocampus.

While earlier behavioral and pharmacological studies on emotional memories utilized narratives as stimuli^1,2,31^, neuroimaging studies have primarily focused on the encoding and recall of static stimuli such as emotionally arousing images^19,20,24,25^ or words^22,23^. Consequently, less is known about the relationship between neural activity and the emotional enhancement of narrative memories. Unlike static stimuli, narratives represent complex experiences where interconnected events unfold dynamically, requiring the accumulation and updating of information over time. Advances in neuroimaging techniques and natural language processing now allow for the study of how the brain encodes and recalls these naturalistic episodes^52^. Here, we take advantage of these advances to investigate the neural basis underlying how emotional arousal enhances what is remembered from temporally extended narratives.

We utilized two publicly available fMRI datasets where participants watched dynamic audiovisual narratives and verbally recalled what they remembered. Arousal ratings of the movies were obtained by analyzing annotations of videos using large language models and from subjective ratings collected in separate behavioral experiments. Applying graph theoretic analyses to the fMRI data, we computed a dynamic measure of functional integration during narrative perception. We assessed the fidelity of participants’ memory by using text embedding models to measure the similarity between their recall and detailed descriptions of the corresponding event. We then tested whether functional integration during encoding mediated the effects of emotional arousal on subsequent recall fidelity. Finally, we replicated our results in a third dataset where we measured arousal using pupillometry while participants listened to and recalled a suspenseful audio story, providing converging evidence of our results with a physiological measure of arousal. Altogether, our work advances an integrative view of arousal-dependent memory enhancement of narrative memories that bridges affective states and ongoing cognition through brain network dynamics.

## Results

Our first set of analyses utilized two publicly available fMRI datasets that, to our knowledge, are the only open datasets combining naturalistic narrative stimuli with free spoken recall. These data allow for fine-grained, event-level assessment of memory fidelity within participants. In the first dataset (“*Film Festival*”)^53^, participants watched 10 audiovisual video clips (range: 2min 12s–7min 45s, average duration: 4min 32s, total duration: 45min 22s). In the second dataset (“*Sherlock*”)^54^, participants watched a 50-minute segment from an episode of the British TV show Sherlock. In both datasets, participants were then asked to describe what they remembered from the movies in as much detail as they could. Each movie was divided into a set of events defined by major shifts in the narrative, including changes in topic, location, time, and characters (68 events in *Film Festival*, average length=38.4s, SD=18.2s; 48 events in *Sherlock*, average length=57.5s, SD=41.76s).

We first obtained detailed annotations of the movie events as a “ground-truth” description of what occurred in each event. To assess how well participants recalled the movie events, we quantified the semantic similarity between participants’ recall transcripts and these annotations. We used Google’s Universal Sentence Encoder (USE)^55^ to convert the annotation and recall transcripts of each event into numerical vectors in a high-dimensional embedding space (see **Methods**). We then calculated the cosine similarity between the vector encoding the annotation and those encoding a participant’s recall of an event as a measure of recall fidelity^56–58^ (**Figure 1**). A high recall fidelity score indicated that a participant’s recall closely matched the annotation of an event. **Extended Data Figure 1** displays an example event in each of the two datasets with the transcripts and fidelity scores of the recall from two different participants.

**Figure 1.**
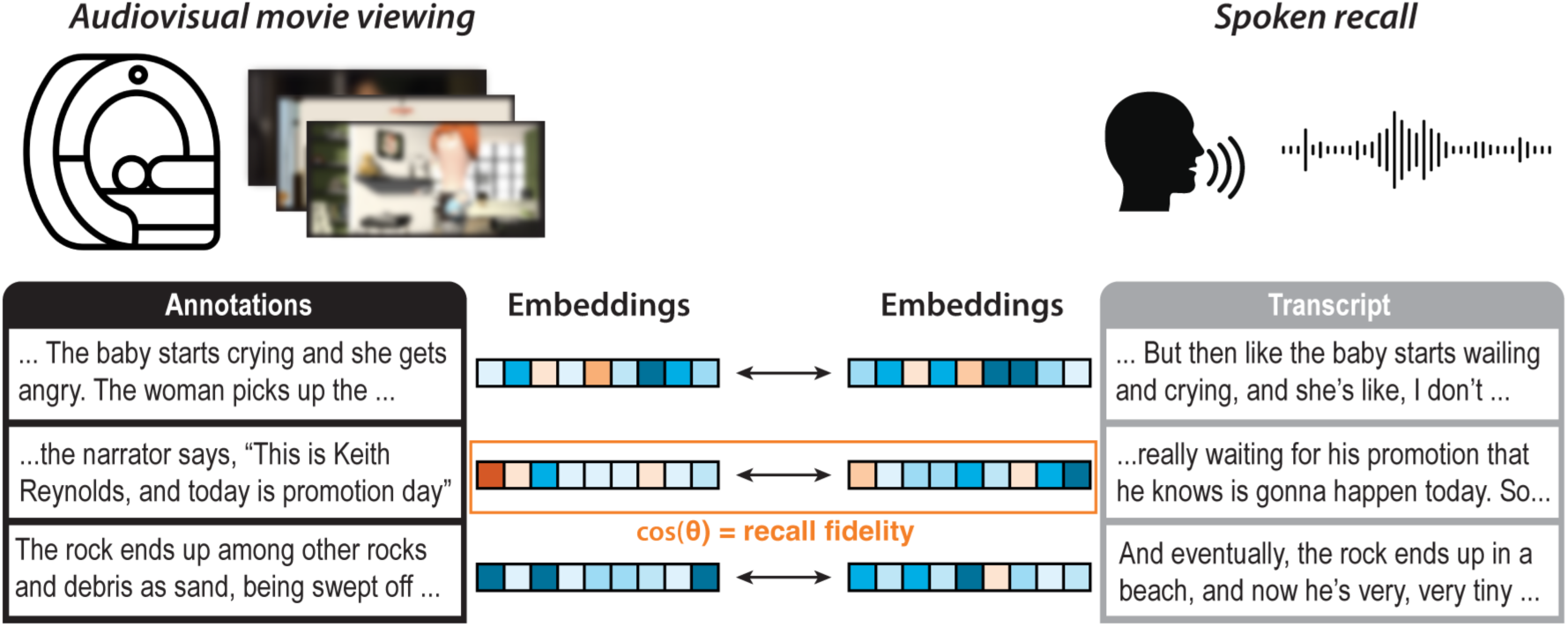
Schematic of task and semantic similarity analysis between movie annotations and spoken recall. Participants watched audiovisual clips in the fMRI scanner and were instructed to recall what they watched from memory. Descriptions of movie events obtained from independent coders and transcriptions of participants’ verbal recall were converted to vector embeddings. Recall fidelity was computed as the cosine similarity between the vector of scene annotations from an event and the transcript from the matching event in a participant’s recall.

### Functional integration during movie-viewing is associated with higher memory fidelity

We first examined how dynamic changes in brain network organization during movie-viewing were associated with subsequent memory fidelity. To that end, we parcellated the brain into 216 cortical and subcortical regions. For each event in each participant, we computed the functional connectivity between each pair of brain regions as the Fisher-*z* transformed Pearson correlation between the BOLD time courses. The connectivity matrices were thresholded to retain the top 15% connections and binarized to create sparse graphs, where nodes represent brain regions and edges represent functional connections between regions. This approach allowed us to leverage graph theory to quantify network properties of large-scale neural dynamics and investigate how functional modules in the brain interact with one another during movie-watching.

In each graph, brain regions were assigned to communities such that regions within a community were more strongly connected to one another than to regions in other communities^59^, thereby partitioning the brain into different functional modules (see **Methods**). We then quantified the diversity of intermodular connections of each brain region by calculating its participation coefficient (*B_T_*)^60^. A brain region with a high *B_T_* has connections distributed across multiple modules, suggesting that it may play a role in integrating information across modules^61^.

Conversely, a brain region with a low *B_T_* has most of its connections within its own module, suggesting more localized processing. High average *B_T_* across brain regions would then reflect high levels of inter-module connectivity, indicating an integrated brain state when there is stronger cohesion across different functional modules.

For each event, we computed the average *B_T_* across brain regions as a measure of whole-brain functional network integration^33^. We note that each graph was thresholded to retain a fixed number of connections, and thus all graphs had the same number of connections. As such, an increase in *B_T_* would not be due to a greater number of connections. Instead, graphs with high average *B_T_* had a greater number of connections *between* functional modules, indicating greater integration across networks (**Figure 2A**). We hypothesized that events associated with greater functional network integration during movie viewing are more strongly encoded and are thus remembered with higher fidelity.

**Figure 2.**
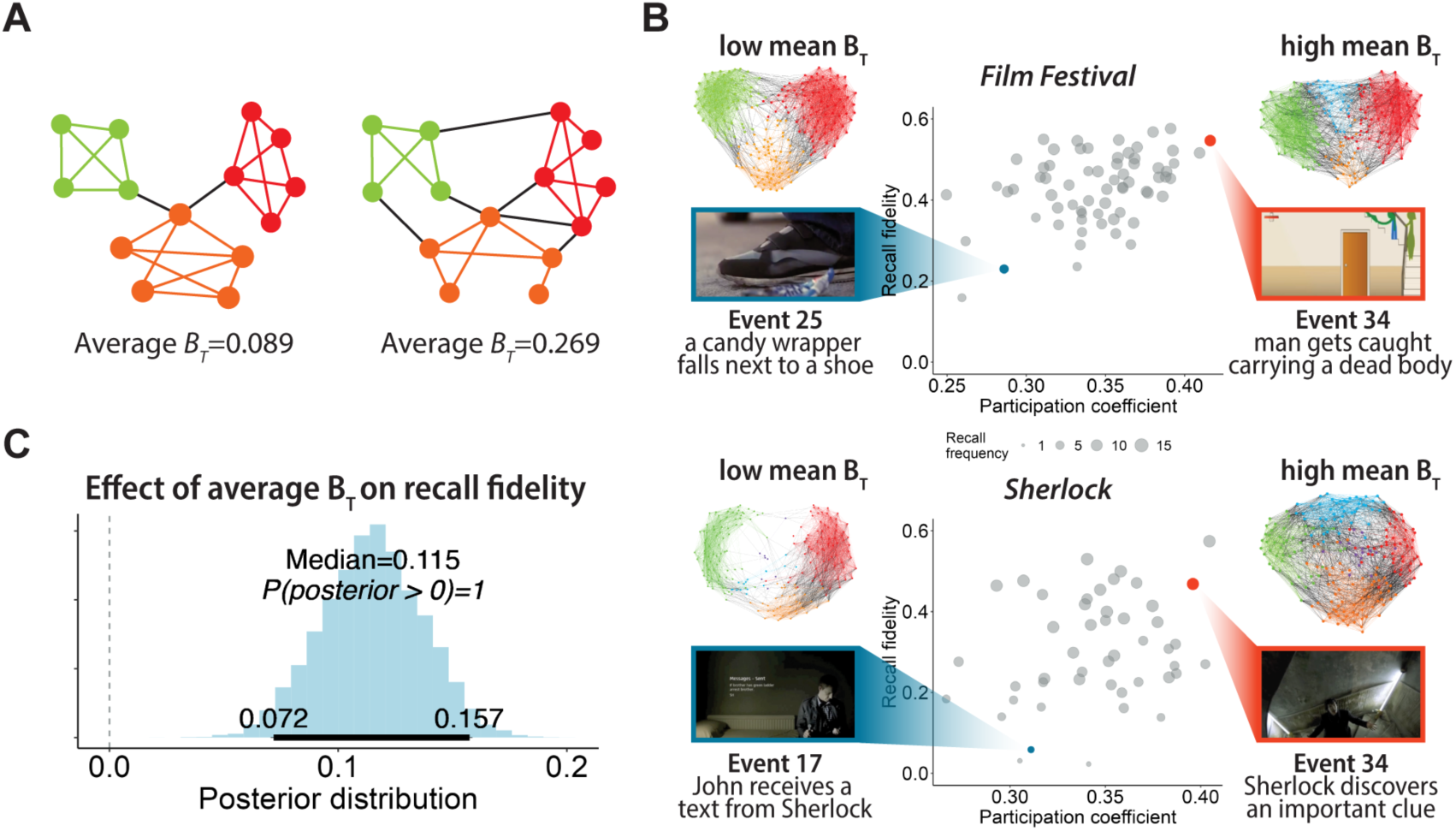
Functional integration during movie viewing is associated with higher recall fidelity. **(A)** Graphical representation of average participation coefficient (*B_T_*). Each community is shown in a different color. Both networks contain the same number of nodes and edges, but the network on the right has more connections between communities, resulting in a higher average *B_T_ .* **(B)** Scatter plots depict the relationship between average *B_T_*, average recall fidelity, and recall frequency. Each datapoint is an event. The size of the circle depicts the number of participants that recalled that event. Red and blue circles correspond to example events with high (red) or low (blue) average *B_T_*. Connectivity plots depict functional connections for the respective events, with colors denoting module assignment. Each connectivity plot contains the same number of connections, but there are more inter-modular connections during events when average *B_T_* is high. **(C)** Posterior distribution of the regression coefficient when predicting recall fidelity from average *B_T_*, estimated by a Bayesian multilevel model that pooled across *Film Festival* and *Sherlock*. The 95% HDI of each distribution is indicated by the bold horizontal line.

We used Bayesian multilevel models to test how functional network integration during movie viewing was related to subsequent event memory. To increase statistical power and estimate reliability, we pooled data across the *Film Festival* and *Sherlock* datasets, treating dataset as a random effect to account for between-dataset variability. Consistent with our hypothesis, average *B_T_* when viewing an event was positively associated with recall fidelity (b=0.11, 95% HDI=[0.07, 0.16], p(b>0)=1, WAIC=4975; **Figure 2B, 2C**). Model comparison showed that this model outperformed a null model without average *B_T_* (WAIC_null_=4996, BF_10_=7739), indicating strong evidence for the contribution of network integration toward recall fidelity.

### Functional integration mediates effects of emotional arousal on narrative recall

Given the well-established role of emotional arousal in enhancing memory encoding^4,17,18^, we next sought to examine the impact of emotional arousal on functional integration and narrative recall. To this aim, we obtained ratings of emotional arousal based on text descriptions of each narrative event using an open-access large-language model (LLM)^62^ (see **Methods**). Additionally, we collected emotional arousal ratings from 30 participants for comparison and validation. Our approach parallels normed affective datasets such as the Affective Norms for English Words^63^ and International Affective Picture System^64^, which use average ratings of arousal and valence across participants to characterize the emotional content of words or images. Similarly, we treated both LLM- and behavior-derived ratings as normative, event-level estimates of emotional arousal of the narrative content.

Human behavioral ratings (*Film Festival*: median=3.2, range=1.3-4.7; *Sherlock*: median=3, range=1.5-4.7) were highly correlated across participants (*Film Festival*: one-to-average *r*=.72, p<.01; Sherlock: one-to-average *r*=.69, p<.01), indicating consistency across individuals on the level of emotional arousal of each event. LLM-generated ratings (*Film Festival*: median=3.3, range=1.6-4.9; *Sherlock*: median=3.5, range=1.9-4.5) correlated with the average behavioral ratings, providing convergent validity of the arousal measures (*Film Festival*: *r*=.60, *p*<.001; *Sherlock*: *r*=.75, *p*<.001; **Extended Data Figure 2**). Subsequent analyses yielded similar results for both the LLM-generated and average behavioral ratings. For brevity, we report results using the LLM-generated ratings here, with results based on behavioral ratings reported in the supplemental materials (*Robustness checks - Behavioral arousal ratings*).

We next sought to relate the arousal ratings to our graph measures of network integration. A Bayesian multilevel model pooling across *Film Festival* and *Sherlock* indicated that average *B_T_* was higher when participants viewed events rated as more emotionally arousing (b=0.12, 95% HDI=[0.07, 0.16], p(b>0)=1, WAIC=5078, WAIC_null_=5105, BF_10_=4.6×10^4^; **Figure 3A**; **Extended Data Figure 3A**), and that events rated as more emotionally arousing were recalled with greater fidelity (b=0.14, 95% HDI=[0.1, 0.18], p(b>0)=1, WAIC=4955, WAIC_null_=4996, BF_10_=3.4×10^7^; **Figure 3B**; **Extended Data Figure 3B**).

**Figure 3.**
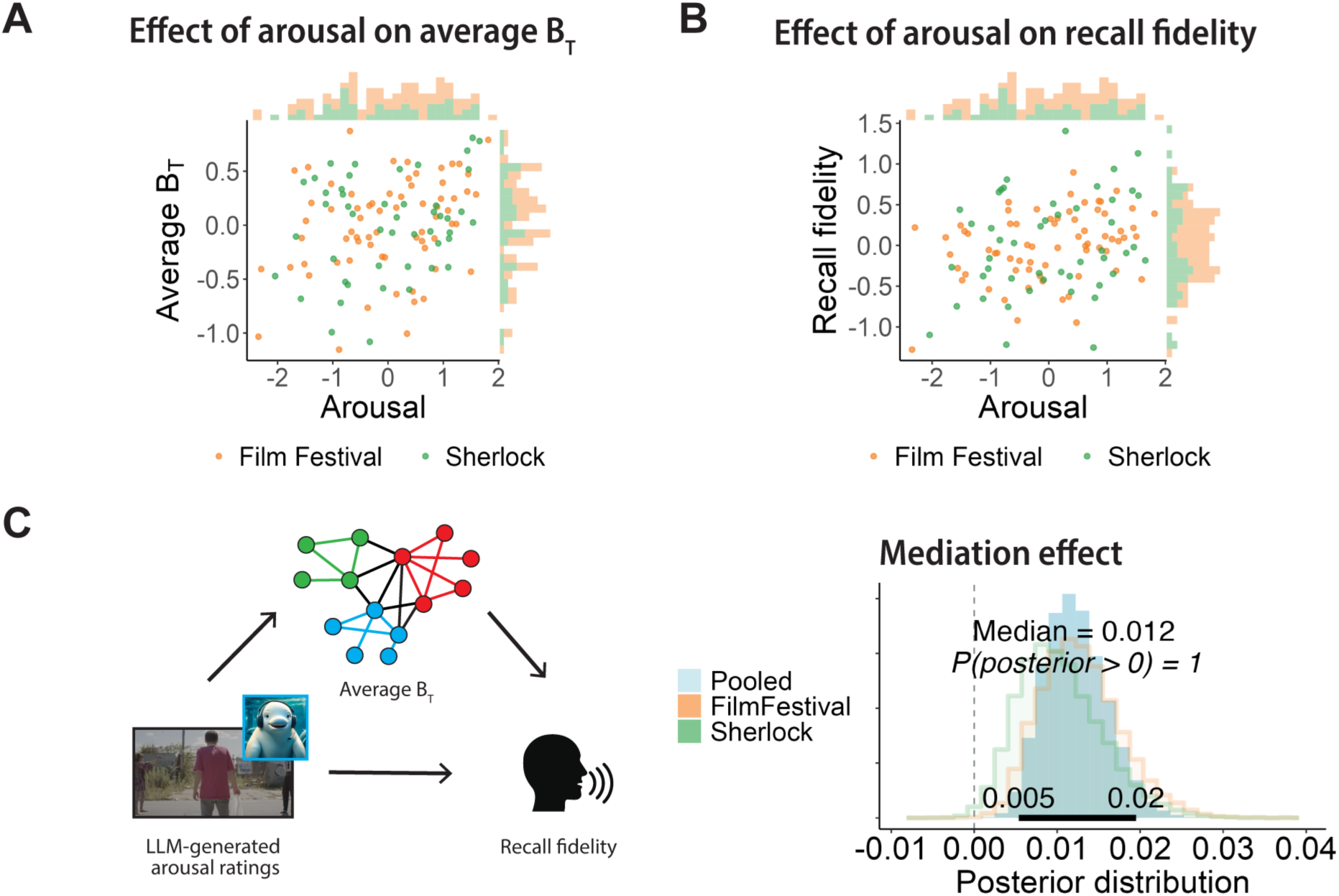
Functional network integration mediates the effects of emotional arousal on recall fidelity. Relationship between emotional arousal and **(A)** average participation coefficient (*B_T_*) and **(B)** recall fidelity. All variables were z-scored across participants. Each point corresponds to an event, with datasets differentiated by color. Marginal histograms display the distribution of arousal and average *B_T_* for each dataset. Corresponding posterior distributions are shown in Extended Data Figure 3. **(C)** Mediation analysis testing whether average *B_T_* during an event mediates the effect of emotional arousal on recall fidelity. The overlaid histograms show the posterior distributions of the mediation (indirect) effect from Bayesian multilevel models estimated separately for each dataset and for the pooled model. The 95% HDI of the regression coefficient estimated using the pooled model is indicated by the bold horizontal line.

Furthermore, average *B_T_* while viewing an event mediated the effects of emotional arousal on recall fidelity (b=0.01, 95% HDI=[0.01, 0.02], p(b>0)=1, WAIC=4943; **Figure 3C**), which accounted for 9% of the total effect of arousal on recall fidelity. Here, we do not compare against a null model, as removing the mediator tests the unique variance it explains in the outcome rather than the presence of mediation. Instead, we rely on the proportion of the posterior distribution of the indirect effect that is greater than zero to assess the strength of evidence for the mediation^65^. To further assess robustness of our results, we show that mediation effect replicates with global efficiency, an alternative measure of whole-brain functional integration (b=0.02, 95% HDI=[0.01,0.03], p(b>0)=1, WAIC=4918).

When event duration, visual intensity, and audio intensity were included as covariates, the mediation effect was attenuated but remained statistically credible, indicating that network integration explained unique variance beyond these features (average *B_T_*: b=0.004, 95% HDI=[0, 0.01], p(b>0)=0.989, WAIC=4833; global efficiency: b=0.005, 95% HDI=[0, 0.01], p(b>0)=0.994, WAIC=4832). Additionally, the mediation effect replicates independently in each dataset, with functional connectivity matrices computed after global signal regression, different parcellation schemes, and network thresholding parameters (see *Robustness checks* in **Supplemental Materials**). Altogether, these findings provide consistent evidence in support of our core hypothesis that emotional arousal enhances memory encoding by facilitating increased functional integration across brain networks.

### Network efficiency within and between canonical brain networks mediates effects of emotional arousal on narrative recall

While average *B_T_* provides a measure of the overall levels of network integration across functional modules in the brain, it is agnostic to the brain regions that drive this integration. Thus, different configurations of large-scale network organization might underlie the observed effects. Are there specific network interactions that mediate the effects of emotional arousal on memory encoding? To address this question, we ran additional analyses to characterize the contributions of connectivity patterns within and between canonical functional brain networks.

We grouped the brain regions based on their assignment to seven functional networks defined by the Schaefer atlas (dorsal attention, DAN; ventral attention, VAN; control, CON; default mode, DMN; visual, VIS; motor, MOT; limbic, LIM), with the 16 subcortical ROIs grouped together as one network (SUB). For each network, we calculated the *within-network efficiency*^66^. For each pair of networks, we calculated the *between-network efficiency*^49^ (see **Methods**). The within- and between-network efficiency provided measures that are often interpreted as reflecting the potential efficiency of information transfer within and across different brain networks, respectively^59,66^.

We tested whether within- and between-network efficiency mediated the effects of emotional arousal on recall fidelity. To correct for multiple comparisons, we thresholded the results at an expected Posterior Error Probability (PEP) of 0.01^67^ (see **Methods**). Within-network efficiency in all 8 functional networks and between-network efficiency in 14 out of 28 possible network pairs mediated the effects of emotional arousal on recall fidelity (**Figure 4A, 4B**; see **Extended Data Table 1** for full statistical detail). Notably, the between-network connections included connections across all 8 functional networks, suggesting that the enhanced encoding of emotionally arousing experiences is supported by brain-wide integration.

**Figure 4.**
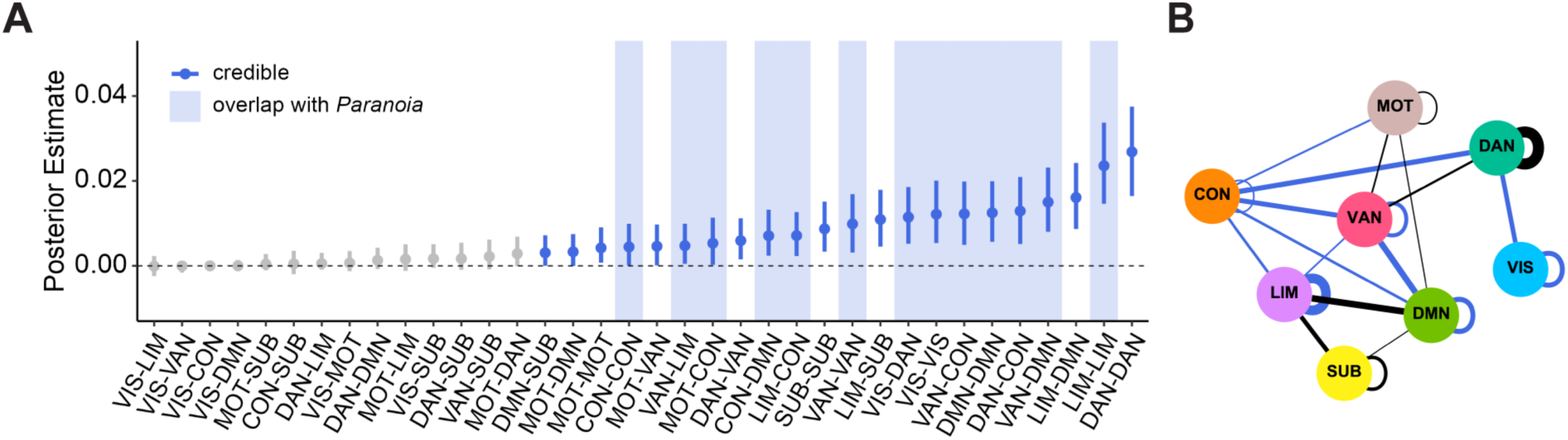
Within- and between-network efficiency mediates the effects of emotional arousal on recall fidelity in audiovisual datasets. **(A)** Dots and whiskers indicate the median and 95% HDI of the posterior distribution of the mediation effect for each network connection. Datapoints are presented in blue if the mediation effect was credible after controlling for multiple comparisons (expected PEP<0.01). Blue shading indicates network connections that were also identified in the *Paranoia* dataset. Network connections were ordered by the median. **(B)** Each node represents a functional network: VIS-visual network, MOT-somatomotor network, DAN-dorsal attention network, VAN-ventral attention network, LIM-limbic network, CON-control network, DMN-default mode network, SUB-subcortical network. Edges represent the credible connections after controlling for multiple comparisons, and edge weights denote the median of the posterior estimate. Black lines indicate edges that were identified only in the audiovisual datasets, while blue edges indicate connections that were also identified in the *Paranoia* dataset.

### Amygdala and hippocampus engagement during emotionally arousing events

Given the significant contributions of individual brain regions to memory and emotional processing, we focused our subsequent analyses on two specific regions of interest commonly implicated in enhanced memory encoding^21,27^: the amygdala and the hippocampus. The events in our study spanned 30 seconds to over a minute (*Film Festival* mean=38.4s, *Sherlock* mean=57.5s). As averaging amygdala and hippocampal activity over the course of an event fails to capture the time-locked neural responses evoked by dynamic stimuli, we employed intersubject correlation (ISC) analyses^68^ to identify shared, stimulus-locked neural responses across participants. Notably, prior work using the *Sherlock* dataset had found that events with higher hippocampal ISC during encoding were more likely to be remembered^54^. Here, we calculated the ISC of the amygdala and hippocampus for each event, and tested their relationship with arousal, integration, and recall fidelity using Bayesian multilevel models that pooled data across both datasets (**Figure 5A, 5B**).

**Figure 5.**
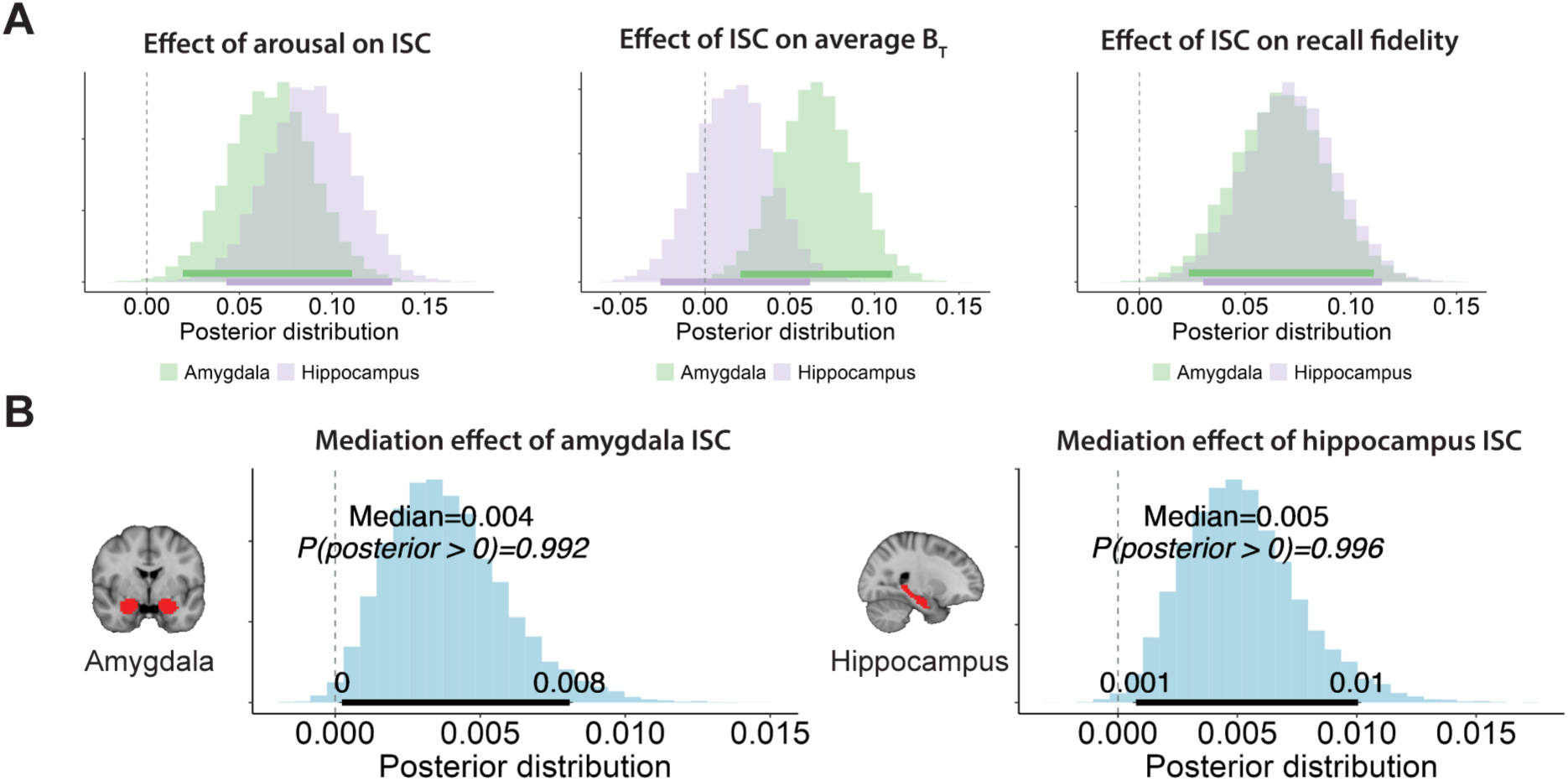
Emotional arousal was associated with hippocampal and amygdalar engagement during encoding. **(A)** Emotionally arousing events were associated with greater inter-subject correlation (ISC) in the amygdala and hippocampus. The overlaid histograms show the posterior distributions from each model for both brain regions, with the 95% HDI in bold horizontal line. In the amygdala, greater ISC was associated with increased arousal (left), stronger integration across functional modules (middle), and higher recall fidelity (right). In the hippocampus, greater ISC was linked to increased arousal and recall fidelity, but not integration. **(B)** Mediation analyses testing whether ISC in the amygdala (left) and hippocampus (right) mediates the effect of emotional arousal on recall fidelity. The 95% HDI of the posterior distribution is indicated by the bold horizontal line.

Amygdala and hippocampal ISC were positively associated with emotional arousal, consistent with the engagement of these regions by emotionally arousing content (amygdala: b=0.07, 95% HDI=[0.02, 0.11], p(b>0)=0.998, WAIC=5197, WAIC_null_=5203, BF_10_=1.5; hippocampus: b=0.09, 95% HDI=[0.04, 0.13], p(b>0)=1, WAIC=5193, WAIC_null_=5206, BF_10_=33). Amygdala ISC was positively associated with average *B_T_* (b=0.07, 95% HDI=[0.02, 0.11], p(b>0)=0.998, WAIC=5094, WAIC_null_=5105, BF_10_=13), while hippocampal ISC was not (b=0.02, 95% HDI=[- 0.03, 0.06], p(b>0)=0.78, WAIC=5102, WAIC_null_=5106, BF_10_=0.3).

Both amygdala and hippocampal ISC were positively associated with recall fidelity (amygdala: b=0.07, 95% HDI=[0.02, 0.11], p(b>0)=0.999, WAIC=4987, WAIC_null_=4996, BF_10_=4.5; hippocampus: b=0.07, 95% HDI=[0.03, 0.11], p(b>0)=1, WAIC=4986, WAIC_null_=4996, BF_10_=10). Additionally, amygdala and hippocampal ISC mediated the effects of arousal on recall fidelity (amygdala: b=0.004, 95% HDI=[0, 0.01], p(b>0)=0.992, WAIC=4950, 3% of total effect; hippocampus: b=0.004, 95% HDI=[0, 0.01], p(b>0)=0.996, WAIC=4948, 4% of total effect). Although inference was based on the pooled model to maximize statistical power, results from individual datasets were directionally consistent in all cases. Most, but not all, dataset-specific estimates met the predetermined credibility threshold of 0.95 (see *Dataset-specific ISC results* in **Supplemental Materials**).

These results are consistent with prior findings indicating the importance of the amygdala and hippocampus to the enhancement of emotional memories. Importantly, the mediation effect of network integration on the relationship between emotional arousal and recall fidelity was robust to controlling for amygdala and hippocampal ISC (b=0.01, 95% HDI=[0, 0.02], p(b>0)=1, WAIC=4933), as well as average activity in the amygdala, hippocampus, and amygdala-hippocampus functional connectivity (b=0.01, 95% HDI=[0.01, 0.02], p(b>0)=1, WAIC=4937), indicating a unique contribution of large-scale functional integration above and beyond that explained by amygdala and hippocampal engagement.

### Functional integration mediates effects of pupil dilation on narrative recall

Thus far, we have demonstrated that emotional arousal, as assessed through subjective ratings and large-language model-derived measures, is linked to heightened network integration and enhanced recall fidelity. Next, we assessed whether our findings would generalize to a physiological measure of arousal. We collected a new dataset (*n*=27) where participants listened to a suspenseful auditory story while their pupil size was continuously measured as a physiological measure of arousal. We further utilized an open dataset (*Paranoia*, *n*=22) of participants listening to the same story while undergoing fMRI^69^. We chose an audio-only stimulus to eliminate luminance-related confounds during pupillometry and to isolate the effects of narrative-driven arousal. Emotional intensity fluctuates over the course of the story^57^, making it well-suited for examining how moment-to-moment changes in arousal relate to event-level memory encoding. Furthermore, the story’s moderate duration (∼20 min) made it feasible to include both narrative listening and free recall in a single session.

The story was segmented into 24 events, and pupil size was averaged for each event. Pupil size across events was correlated between participants (one-to-average *r*=0.6, *p*<0.01), indicating that the story evoked arousal in a reliable manner across participants. Pupil size was associated with greater recall fidelity (b=0.13, 95% HDI=[0.05, 0.2], p(b>0)=0.999, WAIC=1794, BF_10_=15.34; **Figure 6A**), indicating that arousing events were better remembered. Additionally, pupil size was positively associated with greater functional network integration (b=0.33, 95% HDI=[0.26, 0.4], p(b>0)=1, WAIC=1743, BF_10_=94×10^13^; **Figure 6B**). Consistent with our earlier results, functional network integration at encoding was associated with subsequent recall fidelity (b=0.21, 95% HDI=[0.13, 0.28], p(b>0)=1, WAIC=1776, BF_10_=11×10^4^; **Figure 6C**).

**Figure 6.**
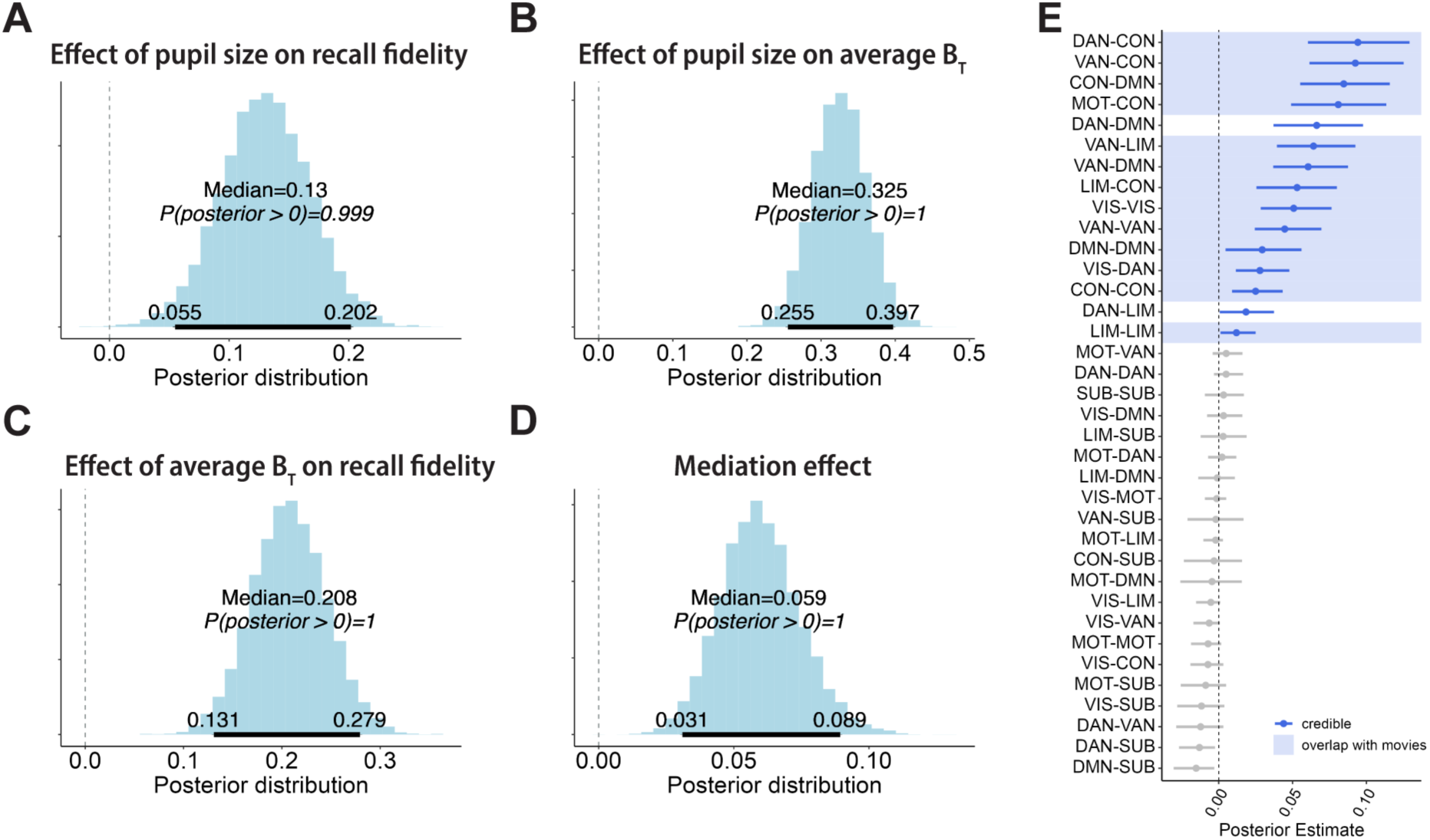
Functional network integration mediates effects of pupil dilation on recall fidelity. Pupil size during story listening was associated with **(A)** recall fidelity and **(B)** average participation coefficient (*B_T_*). **(C)** Average *B_T_* during story listening was associated with recall fidelity, and **(D)** mediated the effects of pupil size on recall fidelity. Posterior distribution was estimated by a Bayesian multilevel model. The 95% HDI of each distribution is indicated by the bold horizontal line. **(E)** Dots and whiskers indicate the median and 95% HDI of the posterior distribution of the mediation effect for each network connection. Datapoints are presented in blue if the mediation effect was credible after controlling for multiple comparisons (expected PEP<0.01). Blue shading indicates network connections that were also identified in the audiovisual datasets (*Film Festival* and *Sherlock*). Network connections were ordered by the median.

Finally, we found that functional network integration mediated the effects of pupil dilation on recall fidelity (b=0.06, 95% HDI=[0.03, 0.09], p(b>0)=1, WAIC=1775; **Figure 6D**), which accounted for 46% of the total effect of arousal on recall fidelity. A post-hoc robustness analysis confirmed that the mediation effect was reliable at the current sample size (**Extended Data Figure 4**). The mediation effect replicated when using global efficiency as the measure of network integration (b=0.09, 95% HDI=[0.06, 0.12], p(b>0)=1, WAIC=1762), and was robust to controlling for event duration and audio intensity (average *B_T_*: b=0.05, 95% HDI=[0.02, 0.07], p(b>0)=1, WAIC=1690; global efficiency: b=0.06, 95% HDI=[0.03, 0.09], p(b>0)=1, WAIC=1685). These results demonstrate that our earlier findings generalize to a physiological measure of arousal.

Within-network efficiency in 5 out of the 8 functional networks (VAN, CON, DMN, VIS, LIM) and between-network efficiency in 10 out of 28 possible network pairs mediated the effects of emotional arousal on recall fidelity (**Figure 6E**; see **Extended Data Table 2** for full statistical detail). Of these 15 connections, 13 overlap with those that were identified in the pooled analysis of the *Film Festival* and *Sherlock* datasets (**Figure 7**). We note that differences between *Paranoia* and the audiovisual datasets should be interpreted with caution as they may be related to lower statistical power in the smaller *Paranoia* sample. Nevertheless, an overlap in the majority of connections across datasets suggest that there is a core set of network interactions supporting the effects of arousal on memory encoding.

**Figure 7.**
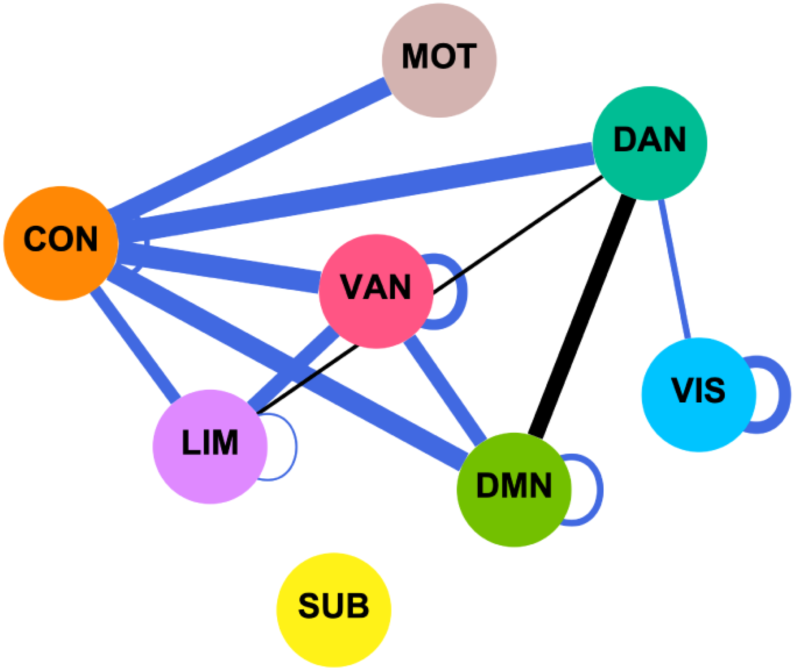
Graphical representation of credible within- and between-network pairs that mediated the effects of pupil size on recall fidelity. Each node represents a functional network: VIS-visual network, MOT-somatomotor network, DAN-dorsal attention network, VAN-ventral attention network, LIM-limbic network, CON-control network, DMN-default mode network, SUB-subcortical network. Edges represent the credible connections after controlling for multiple comparisons, and edge weights denote the median of the posterior estimate. Black lines indicate edges that were identified only in the *Paranoia* dataset, while blue edges indicate connections that were also identified in the pooled audiovisual dataset.

## Discussion

Emotionally arousing experiences are often remembered with great clarity and detail^1,2^. In the present study, we combined fMRI, graph theory, and a naturalistic memory paradigm to study the emotional enhancement of narrative memories. Across three independent fMRI datasets, we demonstrated that whole-brain functional network integration during narrative perception mediated the effects of heightened emotional arousal on the fidelity at which participants subsequently remembered an event. Results replicated across measures of arousal obtained from text analyses of the content, behavioral ratings, and pupil dilation, as well as across movies and audio stories. The mediation effect was observed in interactions across multiple large-scale brain networks, indicating that emotional arousal may enhance memory by promoting coordinated activity across diverse brain regions. Together, our findings suggest that functional network integration supports the enhanced encoding of emotional memories by facilitating coordinated activity across the brain.

Our findings extend the neuroscientific investigation of emotional memories to complex, temporally-extended narratives, which are fundamental to how people make sense of experiences, share knowledge, and connect with others^52,70,71^. Prior work has demonstrated that emotionally arousing content enhances the encoding of narrative memories^1,2,31^. Here, we show that this memory enhancement is associated with increased network integration during narrative encoding. Encoding narratives involves the processing of multisensory information^72^, integration with existing knowledge and schemas^73^, and alignment with ongoing goals and internal states^74^. An integrated brain state, where different functional modules are highly interconnected, facilitates information exchange and coordination across regions^36,75,76^. The enhanced connectivity across perceptual, associative, and memory-related regions may support the binding of narrative events into a coherent memory representation, allowing for better encoding of event details and their broader context.

The current results contribute to the growing body of literature relating arousal to brain network organization. Our findings are consistent with previous studies that demonstrated changes in network organization following shifts in arousal states. For example, Kinnison and colleagues^47^ found increased functional integration across the brain while participants viewed cues indicating potential threats (e.g., a shock) or monetary rewards. Similarly, electrophysiological studies in animal models and fMRI studies in humans have revealed that fluctuations in autonomic arousal induce changes in functional network topology^33,41,43,45,77^. Building upon these findings, our study demonstrates the behavioral consequences of these arousal-dependent network dynamics. Specifically, we show that arousal may prioritize information for encoding by facilitating communication across different functional brain networks, leading to greater fidelity during subsequent recall. These findings highlight the potential importance of arousal-dependent network interactions in shaping ongoing cognition.

One of the strengths of graph theory is its ability to extract network-level metrics that provide insights into how the brain functions as an interconnected system. This approach allows us to capture emergent properties, such as the level of integration across functional modules, that only become apparent when considering the entire network. As a network-level property, integration can arise from various configurations of interactions between functional modules. By framing these network interactions within the broader concept of integration, our approach provides a unifying framework to explain the diverse neural pathways that strengthen emotional memories. Within this framework, we can also examine the specific within and between-network interactions that reliably support emotional memory across contexts, identifying common network configurations that facilitate the encoding of emotionally significant information.

To that end, we identified a core set of network interactions that consistently mediated the effects of emotional arousal on memory. These include interactions between the control network and the ventral attention network (i.e., “salience” network^78^), consistent with prior work showing increased connectivity between these networks during emotionally arousing experiences^42^. The ventral attention network supports the detection of salient information^79^, and increased efficiency between these networks may enhance the allocation of cognitive resources to emotionally significant narrative moments. We also observed consistent mediation effects involving the default mode and the ventral attention networks, which may facilitate the incorporation of externally emotionally salient information into internally constructed narrative models^80,81^. Additionally, mediation by the efficiency between the limbic network and both control and ventral attention networks suggest that arousal-related memory enhancement may depend on coordination between affective processing systems and networks supporting executive control and attentional allocation^82^.

Arousal-related memory enhancement was also mediated by the efficiency between the dorsal attention and control networks. The involvement of the dorsal attention network, which is typically associated with the top-down control of attention, aligns with prior evidence that the enhanced encoding of emotional stimuli depends on attentional processes^83^. From this perspective, attention and arousal are interdependent processes that work in concert to prioritize the encoding of emotionally significant narrative moments. This attentional modulation is potentially supported by the locus coeruleus-norepinephrine (LC-NE) system, which is activated by arousing stimuli, and is known to enhance network integration^33,44,84^ and attention control^85,86^. Consistent with this account, our study found that pupil dilation, a correlate of LC-NE activity^51^, was associated with both network integration and enhanced memory encoding.

The observed mediation effect between the somatomotor and control networks aligns with a recent meta-analysis identifying the pre-supplementary motor area, situated at the intersection of these networks, as one of two regions consistently associated with arousal states (the other being anterior insula)^87^. We also found that efficiency within the visual network and between the visual and dorsal attention networks mediated the effects of arousal on memory, even in *Paranoia*, an audio-only story. Prior work has shown that listening to stories activates early visual areas^88^ and that visual cortex activity encodes schema information from the stories^74^. These findings are often interpreted as evidence of mental imagery during narrative listening. Thus, one interpretation of the involvement of the visual cortex in *Paranoia* is that arousing moments elicit vivid imagery and are selectively prioritized by top-down attention, thereby enhancing memory encoding. Future studies using decoding methods can directly test this account.

While our findings emphasize the importance of network dynamics in enhancing memory encoding, it is equally important to consider the role of specific brain regions within these larger networks. The amygdala and hippocampus, for instance, have been extensively studied for their roles in enhancing emotional memories^19–22,24–27^. Here, we found that emotional arousal engages the amygdala and hippocampus in a stimulus-locked manner. While the amygdala engagement was positively associated with network integration, hippocampal engagement was not, suggesting distinct roles for these regions in coordinating arousal-dependent large-scale network dynamics. Both amygdala and hippocampal engagement were positively associated with recall fidelity and mediated the effects of arousal on memory encoding, extending prior work on images, words, and sounds^19–27^ to temporally extended narratives. Importantly, the mediation effects of network integration remained significant when controlling for amygdala and hippocampal engagement, indicating that the effects of emotional arousal on memory encoding cannot be fully explained by activity in these regions alone.

One potential concern about our design is that the narrative events may not have been sufficiently arousing to engage neural processes implicated in emotional memory enhancement. We consider this unlikely for several reasons. Emotional words have been shown to reliably elicit emotion-related memory effects^22,23^, engaging similar neural processes as highly aversive images. All three narratives in our study contained highly arousing scenes; for example, the *Sherlock* episode included a battlefield sequence and a moment where a main character’s life was under threat - events that are arguably more arousing than isolated words. Indeed, behavioral ratings of emotional arousal in the *Sherlock* and *Film Festival* datasets spanned nearly the full range of the 1–5 scale, suggesting meaningful variability in event-level arousal. Furthermore, heightened arousal was associated with increased amygdala engagement, replicating prior work using images and words. In the *Paranoia* dataset, pupil dilation was synchronized across participants, indicating stimulus-driven fluctuations in arousal that were shared across listeners. These converging behavioral, physiological, and neural measures indicate that the narrative events elicited robust and reliable arousal responses.

Nevertheless, it is important to recognize that watching a movie or listening to a story may differ qualitatively from personally relevant emotional experiences. Extending this work to autobiographical experiences may help clarify how these mechanisms contribute to the formation of vivid and enduring memories formed around personally significant emotional events. This may be particularly relevant in the context of trauma, where emotional memories are often intensely vivid and disruptions in network integration have been observed^89^. Another limitation is that the three datasets were acquired under different conditions and at different sites. While this heterogeneity strengthens the generalizability of our findings, it also complicates cross-dataset comparisons. While sex was relatively balanced across audiovisual and audio datasets, the sample sizes precluded meaningful examination of sex-related differences. We note also that participants were primarily young adults, and future work is needed to assess how these findings extend to older populations, particularly given evidence that LC-linked arousal responses^90^ and neural responses related to event perception undergo age-related changes^91^.

The mediation effects we observed were small in magnitude and were constrained by the modest direct association between emotional arousal and memory fidelity. This is likely due to memory of complex, temporally extended narratives being influenced by many factors beyond emotional arousal, including prior knowledge^92,93^, the extent to which an event aligns with existing schemas^73,94,95^, as well as causal and semantic relationships^53,96,97^. Additionally, our arousal measures were not collected from the fMRI participants, which may have obscured meaningful individual differences in emotional responses. Incorporating participant-specific measures would likely yield stronger direct and mediation effects. Nonetheless, even modest effects can reflect meaningful contributions in naturalistic settings, where memory formation is subject to overlapping influences. Here, we show that mediation via network integration was larger in magnitude than mediation via either hippocampal or amygdala engagement alone. Rather than suggesting that arousal-dependent changes in network integration are the sole driver of memory encoding, our findings identify it as one significant and previously underexplored contributor.

Taken together, our findings provide robust, converging evidence that emotional arousal enhances narrative memories through the functional integration of large-scale brain networks. While our study focused on memory, emotional arousal is also known to influence a broad range of other cognitive processes, including attention^98,99^, perception^100,101^, and decision-making^102,103^. Our work lays the foundation for future research examining interactions operating at different scales, from local activity in the amygdala and hippocampus to brain-wide network dynamics, and how they facilitate the influence of emotional arousal on human cognition and behavior.

## Methods

### Film Festival dataset

Fifteen participants (10 female, 5 male; ages 21 to 33, mean age=27.5) watched 10 short movie clips over the course of two functional runs (TR=1.5s, TE=39ms, flip angle=50°, voxel size=2×2×2mm^3^). Each clip was approximately 5 minutes long (2.15-7.75 min) and varied in content, characters, and genre. Following movie-watching, participants were instructed to verbally describe the clips from memory. Raw anatomical and functional data were downloaded from OpenNeuro^104^. Additional details can be found in the original study^53^. MRI data were preprocessed using FSL/FEAT v.6.00 (FMRIB software library, FMRIB, Oxford, UK). The steps included motion correction of functional images, removal of low-frequency drifts using a temporal high-pass filter (100s cutoff), and spatial smoothing (5-mm FWHM). Functional images were registered to participants’ anatomical image (rigid-body transformation with 6 DOF) and to an MNI template (affine transformation with 12 DOF). Following the original authors of this study^53^, we discarded the first 2 TRs of each run to avoid the non-specific transient increase in BOLD response at the beginning of each run. Data from each run was shifted by 4.5s to account for hemodynamic lag, and then normalized by z-scoring across time within each subject.

### Sherlock dataset

Seventeen participants (7 female, 10 male; ages 19 to 26, mean age=20.8) watched a segment from an episode of the British TV show *Sherlock* over the course of two functional runs (TR=1.5s, TE=28s, voxel size=3×3×4mm^3^). Following movie-watching, participants were instructed to verbally describe the clips from memory. Preprocessed functional and anatomical data were downloaded from Princeton University’s data storage repository, DataSpace^105^. The data had already undergone motion correction, slice timing correction, linear detrending, high-pass filtering, spatial smoothing, normalization to MNI space, z-scoring across each run, and shifting by 4.5s to correct for hemodynamic lag, as detailed in the original publication^54^. No additional preprocessing was applied.

### Paranoia dataset

Twenty-two participants (11 female, 11 male; ages 19 to 35, mean age=27.0) listened to a 20-minute-long story involving a mysterious social event over the course of three functional runs (TR=1s, TE=30ms, flip angle=8°, voxel size=2×2×2mm^3^). Raw anatomical and functional data were downloaded from OpenNeuro^106^. Each run was approximately 7.2 minutes long (5.5-8.7 min). Preprocessing steps (motion correction, smoothing, registration) were identical to those of the *Film Festival* dataset. Data from each run was shifted by 5 TRs to account for hemodynamic lag, and then normalized by z-scoring across time within each subject.

### Movie event segmentation and annotation

For the *Sherlock* stimulus, we utilized the event segmentation and annotation reported in ^54^. Briefly, the 50-minute episode was divided into 48 events (M=57.5s, SD=41.7s) based on significant shifts in the narrative, such as changes in topic, location, time, and characters. An annotator then provided written descriptions about what was happening in the movie during that event. Author K.G. repeated the procedure with the *Film Festival* stimulus, generating 68 events (M=38.4s, SD=18.2s) and corresponding written annotations of each event.

Audio intensity of the *Film Festival, Sherlock,* and *Paranoia* dataset was computed using *audioread* in MATLAB (r2024b, The Mathworks, Natick, MA). Specifically, we extracted the sound envelope by applying the Hilbert transform and taking the absolute value. The envelope signal was then averaged over the event. Frame-wise visual intensity for the *Film Festival* and *Sherlock* datasets was computed by converting each video frame from RGB to HSV color space using MATLAB’s rgb2hsv function. We extracted the “value” (V) channel, which corresponds to brightness, and averaged pixel values within each frame. Visual intensity for each event was then obtained by averaging brightness values across all frames within the event.

### Recall transcripts

Transcripts of verbal memory recall were obtained from ^53^ and ^54^. The transcripts were then segmented and manually matched to the corresponding events based on the annotations that best matched the verbal descriptions. For *Sherlock*, the onset and offset of the events that were remembered were provided by Chen et al. In ^54^. For *Film Festival*, the associated time stamps were identified by author K.G..

### Recall fidelity

We computed the fidelity of participants’ recall by assessing the semantic similarity between participants’ recall transcripts and movie annotations. For each event, we converted both the participants’ recall transcripts and the corresponding movie annotations into separate 512-dimensional vectors using Google’s Universal Sentence Encoder (USE)^55^. Sentences with similar meanings are encoded in vectors that are closer in the embedding space. We chose the USE model because it generates a single embedding vector that captures the overall meaning of text spanning multiple sentences. We then calculated recall fidelity of each event as the cosine similarity between the vector encoding a participant’s recall transcript and that encoding the movie annotation. USE has been frequently employed to measure semantic similarity between event descriptions^38,53,58,107^. Our approach draws on prior work computing the cosine similarity between embedding vectors encoding the annotation of an event and participants’ recall as measures of memory fidelity^56–58^.

### Large language model measure of emotional arousal

We generated ratings of emotional arousal from the event annotations using StableBeluga-13B, (https://huggingface.co/stabilityai/StableBeluga-13B), an open access large language model (LLM)^62^. Researchers are increasingly using LLMs as tools for automated text analysis^108^. For example, LLM-generated valence and arousal ratings of text correspond well to those obtained from behavioral participants^109,110^. We provided the model with event annotations and prompted it to rate the arousal of each event on a scale from 1 to 10. To facilitate comparison with human ratings, which were collected on a 1 to 5 scale, we divided the model-generated ratings by 2. As these ratings were z-scored prior to analyses, the rescaling would not affect the results. As part of the model prompt, we defined arousal as a state of “feeling very mentally or physically alert, activated, and/or energized” (see **Supplemental Materials** for the full model prompt). These instructions were adapted from our earlier work^12^ and drew on dimensional models of affect, where arousal reflects the level of psychological activation or intensity associated with a stimulus or experience, and is orthogonal to valence, which reflects the degree of pleasantness or unpleasantness^10,11^. We avoided using the anchors “calm” and “excited,” as employed in some prior studies^25^, because these terms may be confounded with positive valence and could introduce bias into the LLM’s responses. Due to the inherent stochasticity in the model’s responses, we ran 30 iterations for each event, effectively simulating ratings from 30 “participants”. These ratings were z-scored within each iteration to normalize the data and then averaged across iterations to generate an average arousal rating for each event.

### Behavioral measure of emotional arousal

As a comparison and validation of the LLM-generated arousal ratings, we collected behavioral ratings of emotional arousal from thirty participants (5 male, 24 female, 1 non-binary; ages 21 to 36, mean age=26.23). Experimental procedures were approved by the University of Chicago Institutional Review Board, and participants provided informed consent prior to the start of the study. Participants were compensated with $30 for their time. Participants were provided with the same definition of arousal as the LLM, and were instructed to rate the arousal of each event on a scale of 1 to 5 after watching each event. While the LLM ratings used a 1 to 10 scale to capture finer-grained distinctions, we opted for a coarser 5-point scale for human participants to reduce cognitive load and rating fatigue, given the large number and extended duration of the narrative events. Participants were allowed to pause the video to make their ratings. All participants provided ratings on both *Film Festival* and *Sherlock*, with a short break in between. The ratings were z-scored within each participant and averaged across participants to generate an average arousal rating for each event. We correlated the LLM and behavioral measures to assess the consistency across the two methods of eliciting arousal ratings.

### Generating graphs from functional connectivity matrices

Cortical regions were parcellated into 200 regions of interests (ROI) taken from the Schaefer atlas^111^. Subcortical regions were divided into 16 ROIs based on the Melbourne subcortex atlas^112^. Similar results were observed when repeating our analyses with the Shen parcellation scheme that included 268 parcels encompassing both cortical and subcortical regions (see **Supplementary Material** for additional justification of parcellation scheme). We extracted ROI time courses by averaging all voxels within each ROI. We then regressed out mean white matter and mean cerebrospinal fluid time courses, and framewise displacement from each ROI time course to minimize the influence of motion, non-neuronal noise, and other nuisance signals on the functional connectivity estimates. Global signal regression was not included in the main analyses as it can introduce spurious correlations between brain regions^113,114^. However, we note that our results remained consistent when global signal regression was applied (see **Supplementary Material**).

For each event in each participant, we calculated the Fisher-z transformed Pearson correlation between the BOLD time courses of each pair of ROIs, resulting in a 216×216 functional connectivity (FC) matrix. Each matrix was then thresholded to retain the top 15% strongest connections and binarized to create an unweighted, undirected graph. To ensure the robustness of our findings, we replicated our results at different thresholds (10%, 20% and 25%; see **Supplementary Materials**). All graph theoretic metrics (see below) were then extracted using the Brain Connectivity Toolbox (BCT)^59^.

### Average participation coefficient

Consistent with prior studies^33,35,49^, brain regions were assigned to communities (i.e. modules) using the Louvian algorithm, which seeks to identify a community structure that maximizes the number of within-community connections and minimizes between-community connections. Given the stochastic nature of the algorithm, the community assignment was repeated 1,000 times to obtain a consensus community structure for each participant and event.

We then calculated the participation coefficient (*B_T_*) for each ROI, which quantifies the extent to which the region’s connections are distributed across different communities:

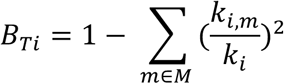

where *k_i_* is the degree (i.e., total number of connections) of region *i*, *k_i,m_* is the number of connections that region *i* has with regions in community *m*. Thus, *B_T_* ranges from 0 to 1 with higher values indicating that a region’s connections are evenly distributed across communities, and lower values indicating that a region’s connections are predominantly within its assigned community. To assess whole-brain network integration, we averaged *B_T_* across all ROIs for each participant and event. The average *B_T_* was used as our primary measure of network integration, with higher values indicating a more integrated brain state during movie-viewing.

### Network efficiency measures

We calculated global efficiency (*Eg*) as an alternative measure of network integration^59^. *Eg* is defined as the average inverse shortest path length between all pairs of nodes in the network:

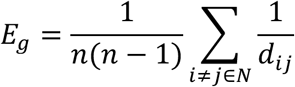

where *n* is the total number of regions, *N* is the set of all regions in the network, and *dij* is the length of the shortest path between nodes *i* and *j*. Higher *Eg* is indicative of greater network integration where fewer connections separate any two brain regions.

Within-network efficiency was calculated as the average inverse shortest path length between all pairs of regions within a network^66^. Between-network efficiency of network 1 and 2 was calculated as the average inverse shortest path length between each region in network 1 and each region in network 2^49^. For all efficiency measures, the shortest path was defined over all regions in the brain. In other words, when calculating between-network efficiency between networks *i* and *j*, the shortest path may include regions outside of the two networks.

### Bayesian multilevel models

We used Bayesian multilevel models to analyze the relationship between brain network metrics, emotional arousal, and recall fidelity. These models were chosen for their ability to account for the hierarchical structure of the data, where events were nested within participants, and to estimate uncertainty in the parameter estimates directly. All models were implemented using the *brms* package (version 2.21.0)^115^ in *R* version 4.3.2, and were run with the random seed “123” to ensure reproducibility. The models included random intercepts for predictors to account for random variability across participants. For models pooling across the Film Festival and Sherlock datasets, we additionally included dataset as a random intercept to account for between-dataset variability.

Models were specified with weakly informative priors to facilitate convergence, while still allowing the posterior distributions to be predominantly shaped by the data (see **Supplementary Materials** for full prior specification). Each model was run with four Markov Chain Monte Carlo (MCMC) chains, each with 4,000 samples, of which the first 1,000 samples were discarded as burn-in. The remaining samples from all chains were concatenated to form the posterior distribution of each parameter. We checked for model convergence by ensuring that the Gelman-Rubin diagnostic (𝑹̂) was less than 1.01 for all parameters. Posterior distributions were summarized by reporting the estimated posterior means and 95% highest density intervals (HDIs)^116^. We considered parameters to be credibly above zero if more than 95% of the posterior distribution was greater than zero.

To evaluate model fit, we computed the Widely Applicable Information Criterion (WAIC) using the *loo* package (version 2.7.0). Lower WAIC values indicate better out-of-sample predictive performance. Where appropriate, we compared models against null models that excluded the predictor of interest. We also computed Bayes Factors (BF_10_) using the *bayestestR* package (version 0.13.2) to quantify evidence relative to the null model. BF_10_ values greater than 1 indicate evidence in favor of the alternative model, with larger values reflecting stronger support.

### Mediation analyses

Mediation effects were computed using the product of coefficients method^117^. Specifically, the posterior distribution of the mediation effect of average *B_T_* was obtained by multiplying the posterior samples of the effect of emotional arousal on average *B_T_* (“path a”) by the posterior samples of the effect of average *B_T_* on recall fidelity while controlling for emotional arousal (“path b”). To evaluate evidence for mediation, we examined the proportion of the posterior distribution of the mediation that was greater than zero. A mediation effect was considered statistically credible if more than 95% of the posterior distribution exceeded zero, and we report the 95% Highest Density Interval (HDI) to describe uncertainty in the estimated effect. For robustness checks, we fit separate models that included event duration, audio intensity, and visual intensity (for audiovisual datasets) as covariates. All other model specifications and inference criteria were identical to the primary models.

We grouped the brain regions into the 7 functional networks defined by the Schaefer atlas (dorsal attention, ventral attention, control, default mode, visual, motor, limbic networks), with the 16 subcortical ROIs grouped together as one network. We then assessed the mediation effect of the within-network efficiency of each network, as well as the between-network efficiency of every pair of networks. To account for multiple comparisons across networks and network pairs, we thresholded the results within each dataset at an expected Posterior Error Probability (PEP) of 0.01, corresponding to a less than 0.01 probability of making an incorrect inference (i.e., concluding that a parameter is greater than 0 when it is in fact less than 0)^67^. From a frequentist perspective, this procedure is analogous to controlling for a false discovery rate at *q* < 0.01^118^. Among the networks and network pairs for which the Highest Density Interval was above zero, we identified those that survived thresholding at an expected PEP < 0.01.

### Intersubject correlation analyses

We extracted the time course data from the amygdala and the hippocampus for each event. For each subject and event, we calculated the inter-subject correlation (ISC) by correlating an individual’s time course data with the average of all other subjects’ data. This provided an ISC value for the amygdala and the hippocampus for each subject and each event. We then tested whether ISC was associated with arousal, memory, and participation coefficient.

### *Paranoia* behavioral dataset

A total of 35 participants were recruited from the University of Chicago research participation pool (SONA systems). The experimental procedures were approved by the Institutional Review Board at the University of Chicago, and all participants provided informed consent at the beginning of the study. Three participants were excluded due to equipment failure; five additional participants were excluded due to noisy pupillometry data (see *Pupillometry* section in **Supplemental Materials**). Thus, 27 participants in total were included in the analysis (12 male, 15 female; ages 20 to 33, mean age=22.3). Detailed procedures regarding noise detection and preprocessing are reported in the supplementary materials.

Pupil size was recorded using the Eyelink 1000 eye tracker (SR Research, Kanata, Ontario, Canada) at a sampling rate of 500 Hz. Prior to recording, participants completed a 5-point calibration sequence. Participants were seated in front of the computer and instructed to fix their eyes on a cross displayed in front of them. After listening to the story, participants were allowed a short break before beginning the recall task. For the recall task, participants were asked to recount the story in their own words. They were encouraged to speak for at least 10 minutes and were permitted to return if they later recalled additional details.

## Data Availability

Raw MRI data from the *Film Festival* dataset are available on OpenNeuro, https://openneuro.org/datasets/ds004042/versions/1.0.1

Preprocessed MRI data from the *Sherlock* dataset are hosted at DataSpace, https://dataspace.princeton.edu/jspui/handle/88435/dsp01nz8062179.

Raw MRI data from the *Paranoia* dataset are available on OpenNeuro, https://openneuro.org/datasets/ds001338/versions/1.0.0

## Code Availability

Custom analysis scripts, as well as behavioral and pupillometry data, are available at: https://github.com/jadynpark/arousal_integration

## Acknowledgements

We thank Drs. Monica Rosenberg and David Gallo for helpful discussion on the work, Drs. Janice Chen and Hongmi Lee for sharing the Film Festival and Sherlock datasets with accompanying annotations. We thank Dr. Emily Finn for sharing the Paranoia dataset. This research was supported by resources provided by the University of Chicago Social Sciences Division, the University of Chicago Research Computing Center and the University of Chicago Data Science Institute.

## Author Contributions Statement

J.S.P. and Y.C.L. conceptualized the analysis. J.S.P., K.G., and J.K. analyzed the data. J.S.P., M.N., I.P., and Y.C.L. wrote the original draft of the manuscript. All authors reviewed and edited the manuscript.

