## Supplementary Information for "Emotional arousal enhances narrative memories through functional integration of large-scale brain networks"

This PDF file includes:

Extended Data Figures 1-4
Extended Data Table 1-2
Supplementary Text
Supplementary References

##### Film Festival

**Event 14: Dinner Table**

The back of a man's head comes into view before the camera pans left to reveal a dinner table with a woman eating at the seat across. The camera cuts back and forth as they cut up their food and eat it. The woman gets up, picks up a bottle of wine, and goes toward the man. She aims to pour him a glass but he covers it with his hand and says, "Not for me dear." He continues to eat. She sits back down at her seat.

**fidelity=0.72**

The segment is subtitled High Maintenance. It opens with a man and a woman sitting at opposite ends of a long dinner table. They're eating in silence. The woman goes to pour the man a glass of wine. He covers the glass with his hand, and says that he cannot drink.

**fidelity=0.28**

But basically it's like a husband and wife together and just like their anniversary. But they don't look very happy, and they're just eating and the wife offers him some wine but he doesn't want to.

##### Sherlock

**Event 7: Get a Cab Discussion**

View of London from the top of a building. October 12th a woman (Helen, a secretary) with a purple shirt and a black skirt is speaking on the phone as she walks across a room in a high rise building. A man's voice (Sir Jeffrey) is heared on the phone saying: "What'd you mean, there's no ruddy car?" Helen, his secretary, replies: "He went to Waterloo. I'm sorry." Sir Jeffrey, a middle-aged man is seen speaking with the woman on his cell phone while walking across the concourse of a busy london railway station. Helen says to him on the phone: "Get a cab". Sir Jeffrey replies while walking: "I never get cabs." Woman looks around to see if anyone is within earshot and says quietly, "I love you," on the phone. Sir Jeffrey asks: "When?" Woman giggles and says "Get a cab!" Sir Jeffrey smiles, gets off the phone and looks around him to get a cab.

**fidelity=0.60**

So then I believe that's when we start to see the first murder. It's like basically, this woman is walking around in her office and she's on the phone with this guy who I guess is her husband. But he's not in the office, he's in a train station, I think. Also on the phone, kind of rushing, and she tells him to get a cab, and he says no, I don't like, or I don't ever use cabs. And then she says OK, but I want to see you soon, or something, go get a cab.

**fidelity=0.37**

Oh but hold on, earlier there might have, I can't remember exactly when these scenes happen, but there are also a series of scenes where we see a few deaths and that presents sort of the plot for the episode, the conflict that SH will eventually have to solve. So the first scene is, we see a man talking with a woman and she says A, you know you should get a taxi and then she says I love you, and B, get a

**Extended Data Figure 1. The semantic similarity between annotations of a movie event and participants' recall transcripts provide a measure of recall fidelity.** Excerpts from the annotations of an example event in each dataset are shown with transcripts and fidelity scores of two different participants recalling that event. Colors indicate matching details between the movie and recall, hand-labeled for illustrative purposes.

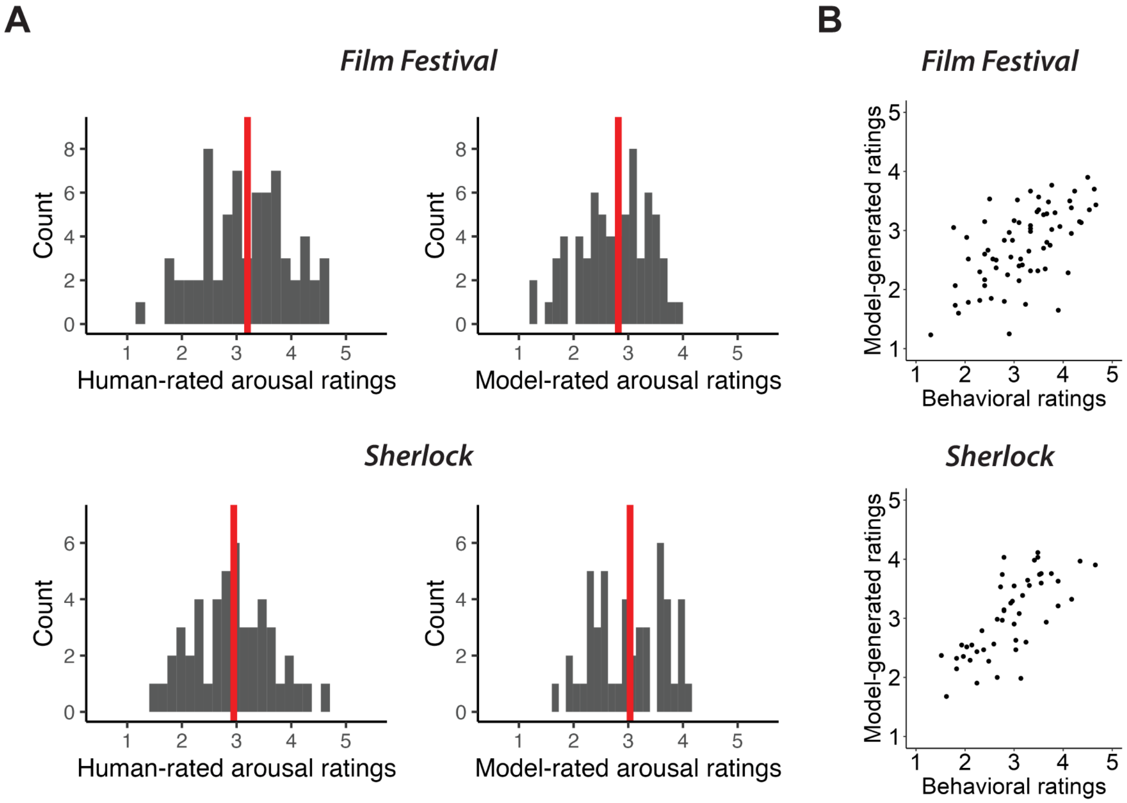


**Extended Data Figure 2. LLM-generated arousal ratings align with human behavioral ratings. (A)** Histogram of human- and model-rated arousal ratings. Arousal ratings for each event were averaged across subjects and iterations (LLM ratings). The red vertical line represents the median. **(B)** Scatterplot of model-arousal ratings against behavioral arousal ratings.

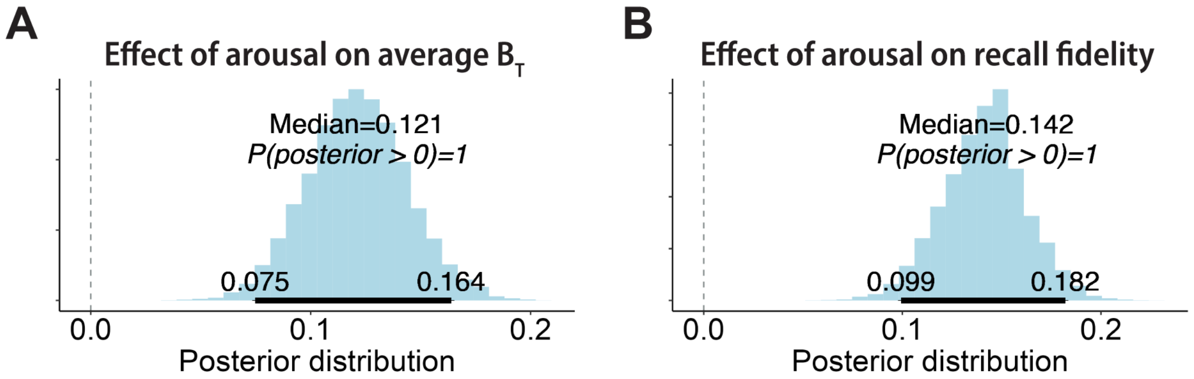


**Extended Data Figure 3. Arousal is associated with functional integration and recall fidelity.** Posterior distribution of regression coefficient when predicting **(A)** average participation coefficient ($B_T$), and **(B)** recall fidelity, estimated by a Bayesian multilevel model that pooled across *Film Festival* and *Sherlock*. The 95% HDI of each distribution is indicated by the bold horizontal line.

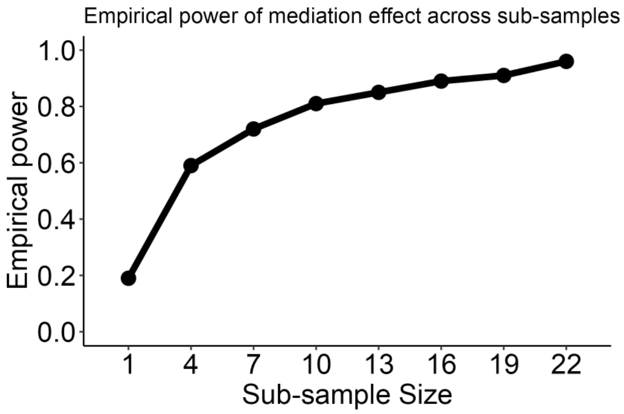


**Extended Data Figure 4. Post-hoc robustness analysis of the mediation effect in the *Paranoia* dataset.** We repeated the mediation analysis with randomly drawn subsets of fMRI participants (with replacement), varying the sub-sample size from 1 to 22. For each subsample size, we conducted 100 iterations and computed empirical power, defined as the proportion of iterations in which more than 95% of the posterior distribution of the mediation effect was greater than zero. Power exceeded 0.8 with as few as 10 participants and approached 1.0 at the full sample size, indicating high reliability of the mediation effect.

**Extended Data Table 1**. Table reporting mediation effect of within- and between-network efficiency on the relationship between emotional arousal and recall fidelity for the audiovisual datasets.

| Network 1 | Network 2 | Median | 95% HDI | p(b>0) | Expected PEP | WAIC |
|---|---|---|---|---|---|---|
| **CON** | **CON** | **0.00** | **[0, 0.01]** | **0.98** | **0.00** | **4,940** |
| **CON** | **DMN** | **0.01** | **[0, 0.01]** | **1.00** | **0.00** | **4,937** |
| CON | SUB | 0.00 | [0, 0] | 0.69 | 0.05 | 4,957 |
| **DAN** | **CON** | **0.01** | **[0.01, 0.02]** | **1.00** | **0.00** | **4,944** |
| **DAN** | **DAN** | **0.03** | **[0.02, 0.04]** | **1.00** | **0.00** | **4,924** |
| DAN | DMN | 0.00 | [0, 0] | 0.92 | 0.01 | 4,954 |
| DAN | LIM | 0.00 | [0, 0] | 0.81 | 0.02 | 4,956 |
| DAN | SUB | 0.00 | [0, 0.01] | 0.90 | 0.01 | 4,955 |
| **DAN** | **VAN** | **0.01** | **[0, 0.01]** | **1.00** | **0.00** | **4,947** |
| **DMN** | **DMN** | **0.01** | **[0.01, 0.02]** | **1.00** | **0.00** | **4,927** |
| **DMN** | **SUB** | **0.00** | **[0, 0.01]** | **0.99** | **0.00** | **4,953** |
| **LIM** | **CON** | **0.01** | **[0, 0.01]** | **1.00** | **0.00** | **4,945** |
| **LIM** | **DMN** | **0.02** | **[0.01, 0.02]** | **1.00** | **0.00** | **4,925** |
| **LIM** | **LIM** | **0.02** | **[0.01, 0.03]** | **1.00** | **0.00** | **4,917** |
| **LIM** | **SUB** | **0.01** | **[0, 0.02]** | **1.00** | **0.00** | **4,941** |
| **MOT** | **CON** | **0.01** | **[0, 0.01]** | **0.98** | **0.00** | **4,953** |
| MOT | DAN | 0.00 | [0, 0.01] | 0.98 | 0.00 | 4,950 |
| **MOT** | **DMN** | **0.00** | **[0, 0.01]** | **0.99** | **0.00** | **4,951** |
| MOT | LIM | 0.00 | [0, 0.01] | 0.88 | 0.02 | 4,956 |
| **MOT** | **MOT** | **0.00** | **[0, 0.01]** | **1.00** | **0.00** | **4,949** |
| MOT | SUB | 0.00 | [0, 0] | 0.76 | 0.04 | 4,957 |
| **MOT** | **VAN** | **0.01** | **[0, 0.01]** | **0.99** | **0.00** | **4,952** |
| **SUB** | **SUB** | **0.01** | **[0, 0.02]** | **1.00** | **0.00** | **4,945** |
| **VAN** | **CON** | **0.01** | **[0, 0.02]** | **1.00** | **0.00** | **4,941** |
| **VAN** | **DMN** | **0.01** | **[0.01, 0.02]** | **1.00** | **0.00** | **4,929** |
| **VAN** | **LIM** | **0.01** | **[0, 0.01]** | **0.99** | **0.00** | **4,953** |
| VAN | SUB | 0.00 | [0, 0.01] | 0.94 | 0.01 | 4,956 |
| **VAN** | **VAN** | **0.01** | **[0, 0.02]** | **1.00** | **0.00** | **4,948** |
| VIS | CON | 0.00 | [0, 0] | 0.52 | 0.07 | 4,958 |
| **VIS** | **DAN** | **0.01** | **[0.01, 0.02]** | **1.00** | **0.00** | **4,933** |
| VIS | DMN | 0.00 | [0, 0] | 0.55 | 0.06 | 4,958 |
| VIS | LIM | -0.00 | [0, 0] | 0.45 | 0.08 | 4,957 |
| VIS | MOT | 0.00 | [0, 0] | 0.80 | 0.03 | 4,953 |
| VIS | SUB | 0.00 | [0, 0.01] | 0.94 | 0.01 | 4,953 |
| VIS | VAN | -0.00 | [0, 0] | 0.44 | 0.10 | 4,958 |
| **VIS** | **VIS** | **0.01** | **[0.01, 0.02]** | **1.00** | **0.00** | **4,921** |

Note: Rows in bold font indicate networks and network pairs where the expected PEP was <0.01 and the 95% HDI were >0. p(b>0): proportion of the posterior draws that were >0. WAIC: Widely Applicable Information Criterion.

**Extended Data Table 2.** Table reporting mediation effect of within- and between-network efficiency on the relationship between pupil dilation and recall fidelity for the *Paranoia* dataset.

| Network 1 | Network 2 | Median | 95% HDI | p(b>0) | Expected PEP | WAIC |
|---|---|---|---|---|---|---|
| **CON** | **CON** | **0.03** | **[0.01, 0.04]** | **1.00** | **0.00** | **1,783** |
| **CON** | **DMN** | **0.09** | **[0.06, 0.12]** | **1.00** | **0.00** | **1,712** |
| CON | SUB | -0.00 | [-0.02, 0.02] | 0.37 | 0.13 | 1,797 |
| **DAN** | **CON** | **0.09** | **[0.06, 0.13]** | **1.00** | **0.00** | **1,757** |
| DAN | DAN | 0.01 | [0, 0.02] | 0.90 | 0.01 | 1,795 |
| **DAN** | **DMN** | **0.07** | **[0.04, 0.1]** | **1.00** | **0.00** | **1,774** |
| **DAN** | **LIM** | **0.02** | **[0, 0.04]** | **0.98** | **0.00** | **1,792** |
| DAN | SUB | -0.01 | [-0.03, 0] | 0.00 | 0.39 | 1,788 |
| DAN | VAN | -0.01 | [-0.03, 0] | 0.05 | 0.29 | 1,794 |
| **DMN** | **DMN** | **0.03** | **[0, 0.06]** | **0.99** | **0.00** | **1,717** |
| DMN | SUB | -0.01 | [-0.03, 0] | 0.00 | 0.37 | 1,790 |
| **LIM** | **CON** | **0.05** | **[0.03, 0.08]** | **1.00** | **0.00** | **1,713** |
| LIM | DMN | -0.00 | [-0.01, 0.01] | 0.43 | 0.09 | 1,779 |
| **LIM** | **LIM** | **0.01** | **[0, 0.03]** | **0.99** | **0.00** | **1,786** |
| LIM | SUB | 0.00 | [-0.01, 0.02] | 0.66 | 0.07 | 1,797 |
| **MOT** | **CON** | **0.08** | **[0.05, 0.11]** | **1.00** | **0.00** | **1,764** |
| MOT | DAN | 0.00 | [-0.01, 0.01] | 0.72 | 0.04 | 1,788 |
| MOT | DMN | -0.01 | [-0.03, 0.02] | 0.33 | 0.15 | 1,797 |
| MOT | LIM | -0.00 | [-0.01, 0] | 0.18 | 0.20 | 1,795 |
| MOT | MOT | -0.01 | [-0.02, 0] | 0.04 | 0.33 | 1,787 |
| MOT | SUB | -0.01 | [-0.03, 0.01] | 0.11 | 0.22 | 1,768 |
| MOT | VAN | 0.01 | [0, 0.02] | 0.88 | 0.01 | 1,795 |
| SUB | SUB | 0.00 | [-0.01, 0.02] | 0.72 | 0.05 | 1,796 |
| **VAN** | **CON** | **0.09** | **[0.06, 0.13]** | **1.00** | **0.00** | **1,741** |
| **VAN** | **DMN** | **0.06** | **[0.04, 0.09]** | **1.00** | **0.00** | **1,752** |
| **VAN** | **LIM** | **0.06** | **[0.04, 0.09]** | **1.00** | **0.00** | **1,745** |
| VAN | SUB | -0.00 | [-0.02, 0.02] | 0.42 | 0.11 | 1,797 |
| **VAN** | **VAN** | **0.04** | **[0.02, 0.07]** | **1.00** | **0.00** | **1,768** |
| VIS | CON | -0.01 | [-0.02, 0] | 0.06 | 0.27 | 1,794 |
| **VIS** | **DAN** | **0.03** | **[0.01, 0.05]** | **1.00** | **0.00** | **1,782** |
| VIS | DMN | 0.00 | [-0.01, 0.02] | 0.74 | 0.03 | 1,797 |
| VIS | LIM | -0.01 | [-0.02, 0] | 0.04 | 0.32 | 1,794 |
| VIS | MOT | -0.00 | [-0.01, 0.01] | 0.28 | 0.18 | 1,792 |
| VIS | SUB | -0.01 | [-0.03, 0] | 0.06 | 0.25 | 1,794 |
| VIS | VAN | -0.01 | [-0.02, 0] | 0.03 | 0.35 | 1,791 |
| **VIS** | **VIS** | **0.05** | **[0.03, 0.08]** | **1.00** | **0.00** | **1,767** |

Rows in bold font indicate the networks and network pairs where the expected PEP was <0.01 and the 95% HDI were >0. p(b>0): proportion of the posterior draws that were >0. WAIC: Widely Applicable Information Criterion.

### Supplementary Text

#### Robustness checks

***Dataset-specific mediation results.*** In the main text, we report results based on data pooled from the *Film Festival* and *Sherlock* datasets. In addition to these analyses, we assessed the robustness and generalizability of our findings by testing each dataset independently. The mediation effect of the average participation coefficient on arousal and recall was observed independently in both datasets (*Film Festival*: b=0.01, 95% HDI=[0, 0.02], p(b>0)=0.999, WAIC=2716; *Sherlock:* b=0.01, 95% HDI=[0, 0.02], p(b>0)=0.995, WAIC=2228), supporting the replicability of the main results across independent samples.

***Alternative parcellation schemes.*** In the main text, we report results from a parcellation scheme that included 200 cortical ROIs from the Schaefer parcellation and 16 subcortical ROIs from the Melbourne subcortical atlas. We chose the Schaefer parcellation scheme because it was derived using both task-based and resting state fMRI data (1). We further chose the parcellation with 200 ROIs because it approximately matched the number of ROIs used in previous work examining the relationship between memory and network integration (2). We then added the subcortical ROIs given the importance of these regions to both emotional processing and memory encoding.

However, we do not think that our results depend on the specific parcellation scheme used. To test the robustness of our findings, we repeated our analysis with the Shen parcellation scheme (3), which consists of 268 ROIs including both cortical and subcortical regions. Results using the Shen parcellation scheme were consistent with those reported in the main text (b=0.01, 95% HDI=[0, 0.02], p(b>0)=0.999, WAIC=4950), indicating that our results were not specific to the choice of parcellation.

***Varying thresholds for FC matrices.*** In our main analysis, we set a proportional threshold ($\tau$) of 0.15. To ensure the robustness of our findings, we tested a range of different threshold values ($\tau$=0.10, 0.20, 0.25), in line with previous literature (2, 4, 5). Across all thresholds, we found a positive association between arousal and average $B_T$, and average $B_T$ and recall fidelity. Average $B_T$ also reliably mediated between arousal and recall fidelity ($\tau$=0.10: b=0.01, 95% HDI=[0, 0.02], p(b>0)=0.999, WAIC=4946; $\tau$=0.20: b=0.01, 95% HDI=[0.01, 0.02], p(b>0)=1, WAIC=4939; $\tau$=0.25: b=0.01, 95% HDI=[0, 0.02], p(b>0)=0.999, WAIC=4944). This suggests that the relationship between arousal, functional network integration, and recall fidelity is robust across various thresholds.

***Global signal regression.*** In the analyses reported in the main text, we did not include global signal regression due to the potential for introducing spurious correlations and anti-correlations between brain regions (6,7). However, when we repeated the mediation analyses with global signal regression, the results remained consistent (b=0.01, 95% HDI=[0, 0.01], p(b>0)=0.999, WAIC=4899), suggesting that our findings were not driven by global noise or artifacts.

***Behavioral arousal ratings.*** Large language models offer a transparent, reproducible, and scalable approach for sentiment analysis (8). Recent work has shown that affective ratings of text generated using LLMs correspond well to those obtained from human participants (9,10). Here, we validated the LLM-generated arousal ratings from the movie annotations against ratings collected from human participants. The mediation results were replicated using the behavioral arousal ratings (b=0.01, 95% HDI=[0, 0.02], p(b>0)=0.999, WAIC=4917), indicating that our findings are robust across different methods of measuring arousal, providing converging evidence for our conclusions.

**Dataset-specific ISC results**

In the main text, we report ISC results from data pooled across the *Film Festival* and *Sherlock* datasets to maximize statistical power. Here, we report findings obtained from each dataset separately. Results from individual datasets were directionally consistent in all cases. Most, but not all, dataset-specific estimates met the predetermined credibility threshold of 0.95.

***Effect of Arousal on ISC***

Amygdala ISC was positively associated with arousal:

- *Film Festival*: b=0.06, 95% HDI=[0.01, 0.13], p(b>0)=0.979
- *Sherlock*: b=0.07, 95% HDI=[0, 0.14], p(b>0)=0.98

Hippocampal ISC also showed a positive association with arousal:

- *Film Festival*: b=0.06, 95% HDI=[0, 0.12], p(b> 0)=0.969
- *Sherlock*: b=0.13, 95% HDI=[0.06, 0.20], p(b>0)=0.999

***Effect of ISC on Average $B_T$***

Amygdala ISC was associated with higher between-network integration ($B_T$):

- *Film Festival*: b=0.07, 95% HDI=[0.02, 0.13], p(b>0)=0.993
- *Sherlock*: b=0.05, 95% HDI = [–0.02, 0.12], p(b > 0)=0.932 (below threshold)

Hippocampal ISC was not associated with Average $B_T$ in either dataset, consistent with results from the pooled model:

- *Film Festival*: b = 0.02, 95% HDI = [–0.04, 0.08], p(b>0)=0.732
- *Sherlock*: b = 0.01, 95% HDI = [–0.05, 0.08], p(b>0)=0.663

***Effect of ISC on Recall Fidelity***

The relationship between ISC and recall fidelity was credible only in *Sherlock* and not *Film Festival*.

Amygdala ISC and recall fidelity:

- *Film Festival*: b=0.03, 95% HDI = [–0.02, 0.09], p(b>0)=0.871 (below threshold)
- *Sherlock*: b=0.11, 95% HDI=[0.05, 0.18], p(b>0)=0.999

Hippocampal ISC and recall fidelity:

- *Film Festival*: b=0.03, 95% HDI=[–0.03, 0.09], p(b>0)=0.815 (below threshold)
- *Sherlock*: b = 0.13, 95% HDI = [0.06, 0.20], p(b>0)=0.999).

##### *Mediation Analyses*

In *Sherlock*, both amygdala and hippocampal ISC mediated the effect of arousal on recall fidelity:

- Amygdala ISC: b=0.01, 95% HDI=[0, 0.02], p(b>0)=0.979
- Hippocampus ISC: b=0.01, 95% HDI=[0, 0.03], p(b>0)=0.999

In *Film Festival*, we did not conduct mediation analyses due to the absence of significant ISC-recall associations.

#### Pupillometry

##### *Preprocessing*

*Exclusion criteria*

We aligned participants' data to stimulus and calculated the difference between every consecutive sample. Large fluctuations between consecutive samples is indicative of noisy data. For each participant, we calculated the proportion of samples that deviated from the median pupil size by 2 standard deviations. Participants were excluded if more than a quarter of the data points fell outside this criterion. Five out of 32 participants were excluded following this procedure.

*Blink and artifact detection*

For the remaining participants, blinks were detected based on Eyelink's algorithm with default settings. The identified blinks were interpolated using the samples immediately preceding and following them. Similarly, any other missing data shorter than a second were interpolated. The interpolated data were then downsampled to the sampling rate of 50 Hz. The downsampled data were segmented into 1-second epochs to align with the TR of the fMRI data, resulting in 50 samples per epoch. We calculated the average pupil size within each epoch. Artifactual samples were identified as those falling above or below 3 standard deviations from the epoch mean. If more than 40% of the samples in an epoch were deemed as artifactual samples, the mean pupil diameter for that epoch was replaced via linear interpolation across adjacent epochs.

#### Large language model (LLM) prompt

StableBeluga-13B is an auto-regressive language model fine-tuned on Llama2 13B. In particular, the model is fine-tuned to recognize and respond to a specific prompt format, which structures the interaction between the user and the model. The structured format helps the model differentiate between components of the input and generate appropriate responses based on user requests. The prompt format consists of three parts: 1. a *System Prompt* that sets the context for the interaction, guiding the model on how to approach the task; 2. a *User Prompt* containing the instructions generated by the user, directing the model on what it needs to do; and 3. an *Assistant Response* where the model generates its output based on the provided instructions.

The model used in this study is available at: https://huggingface.co/stabilityai/StableBeluga-13B, which was downloaded onto the University of Chicago Data Science Institute High Performance Computing Cluster for analyses. The temperature of the model, which determines the variability in the model's response, was set to 0.85. Below is the prompt used to rate each event:

> ### System: You are Stable Beluga 13B, an AI that follows instructions extremely well. Help as much as you can. Remember, be safe, and don't do anything illegal.
>
> ### User: Arousal refers to when you are feeling very mentally or physically alert, activated, and/or energized. Read the following description of a scene and rate the arousal level of the scene on a scale of 1 to 10, with 1 being low arousal and 10 being high arousal. Please give a numeric rating. Only give the rating; no need to provide explanations.
>
> Scene: [Annotation of the movie event]
>
> ### Assistant: [Model-generated arousal rating]

**Priors for Bayesian multilevel models**

For each subject $i$, event $j$, and dataset $k$, the likelihood for an observed outcome $y_{i,j,k}$ is assumed to be drawn from a normal distribution:

$$y_{i,j,k} \sim Normal(\mu_{i,j,k}, \sigma)$$

where $\mu_{i,j}$ denotes the expected outcome for subject $i$, event $j$, and dataset $k$, and $\sigma$ is the residual standard deviation of the outcome.

$\mu_{i,j}$ is modeled as a function of fixed intercept and fixed slope for the predictors with subject- and dataset-specific random intercepts:

$$\mu_{i,j,k} = \beta_0 + \alpha_{\text{subj[i]}} + \alpha_{\text{dataset[k]}} + \beta_1 x_{i,\text{j,k}}$$

$$\beta_0 \sim Normal(0,1)$$

$$\beta_1 \sim Normal(0,1)$$

$$\alpha_{\text{subj[i]}} \sim Normal(0, \sigma_{\text{subj}})$$

$$\alpha_{\text{dataset}[k]} \sim \mathit{Normal}(0, \sigma_{\text{dataset}})$$

$$\sigma_{\text{subj}} \sim \mathit{Exponential}(1)$$

$$\sigma_{\text{dataset}} \sim \mathit{Exponential}(1)$$

$$\sigma \sim \mathit{exponential}(1)$$

These priors were designed to be weakly informative. The Normal(0, 1) priors for fixed effects are centered at zero, reflecting no strong prior belief about the direction of the effect, while allowing plausible values within a reasonable range. The standard deviation of 1 provides light regularization, helping to prevent overfitting without overly constraining the model. The Exponential(1) priors on the standard deviation parameters restrict values to be positive and favor smaller variances without strongly constraining the model.

To confirm that our results were not dependent on our choice of priors, we ran the mediation models using less informative priors. To minimize assumptions about its distribution, we applied uniform priors to the slope parameters; to account for the possibility of extreme values, we used a Cauchy(0, 2) prior for the standard deviation, which has a heavier tail compared to the exponential distribution. The results remained consistent under these prior specifications (audiovisual: b=0.01, 95% HDI=[0, 0.02], p(b>0)=1, WAIC=4956; *Paranoia*: b=0.06, 95% HDI=[0.03, 0.09], p(b>0)=1, WAIC=1775), suggesting that our results are robust to different prior assumptions.
